# Systematic feature and architecture evaluation reveals tokenized learned embeddings enhance siRNA efficacy prediction

**DOI:** 10.1101/2025.08.12.669916

**Authors:** Rory Coffey

## Abstract

Recent advances in machine learning have improved the prediction of siRNA efficacy, with graph neural networks and transformer-based encodings leading the way. However, existing models still face challenges, including potential inaccuracies in thermodynamic feature calculations (such as incorrect strand selection for siRNA-mRNA Gibbs free energy), limited effective utilization of available datasets, and a lack of systematic model refinement. In this study, I systematically evaluated the predictive power of individual features and neural network architectures to identify the most effective configurations. This process led to the development of RN.Ai-Predict, a model built upon a tokenized learned embedding for nucleotide sequences. This work demonstrates that a methodical approach to feature selection and hyperparameter tuning, particularly favoring learned embeddings, can yield a more accurate and reliable model for predicting siRNA efficacy, outperforming more complex architectures in generalizability.

---

Early efforts to predict siRNA efficacy relied on rule-based approaches, which identified specific sequence and thermodynamic properties associated with effective gene silencing (*1–6*). A prominent example is the set of eight criteria developed by Reynolds et al., which remains widely influential:

1. 30%–52% G/C content
2. At least 3 ‘A/U’ bases at positions 15–19
3. Absence of internal repeats
4. An ‘A’ base at position 19
5. An ‘A’ base at position 3
6. A ‘U’ base at position 10
7. A base other than ‘G’ or ‘C’ at 19
8. A base other than ‘G’ at position 13

These rule-based methods established fundamental guidelines for siRNA design but were constrained by their inability to account for complex, nonlinear interactions within sequence data. While certain design rules remained consistent across studies, others diverged, highlighting the need for more sophisticated predictive models.

Machine learning (ML) subsequently offered data-driven alternatives. Initial ML models included linear approaches like i-Score and DSIR (*7, 8*), followed by ensemble methods such as Random Forest (e.g., Monopoli-RF, siRNAPred (*9–11*)) and Support Vector Machines (*12*). These improved upon rule-based systems but were still limited in capturing intricate feature interactions. Deep learning (DL) has further advanced the field, with architectures like Deep Neural Networks (DNNs) (*13*), Convolution Neural Networks (CNNs) for local pattern extraction (*14*), Graph Neural Networks (GNNs) for modeling siRNA-target interactions (*15, 16*), and Transformer models leveraging self-attention for contextual nucleotide relationships (*17*). While powerful, the performance of these models may be highly dependent on evaluation strategies, and their ability to generalize to entirely new gene targets requires rigorous validation.

The performance of these predictive models is critically dependent on their input features, which are generally grouped into two categories: nucleotide sequence representations and siRNA-mRNA interaction thermodynamics. Within the nucleotide category, effective models consider more than just the siRNA sequence itself. Studies have shown that the 19-20 nucleotides of mRNA flanking both sides of the siRNA binding site also significantly influence efficacy (*14, 17*).

For **nucleotide sequence representation** within machine learning and neural network models, two methods are common: one-hot encoding and k-mer counts (*14–17*). One-hot encoding represents each nucleotide as a 4-dimensional binary vector, e.g. G=[0, 0, 1, 0], whereas k-mer counts finds the number of occurrences of all possible tri-nucleotide sequences. An alternative proposed in this study, inspired by natural language processing, is to learn a dense vector representation, also known as an embedding, for each nucleotide (*18, 19*). Instead of being fixed, these embeddings are optimized during model training, allowing the network to place nucleotides in a conceptual vector space based on the properties most predictive of siRNA efficacy. For example, the model might learn to position ‘G’ and ‘C’ closely to represent their contribution to thermodynamic stability, while simultaneously positioning ‘G’ and ‘A’ together when their shared purine structure (e.g., steric hindrance) is more influential. This ability to learn task-specific, latent biochemical and positional features is a key advantage. While large, pre-trained embeddings like those from RNA-FM are being explored (*17, 20*), the power of embeddings learned de novo for the specific task of siRNA efficacy prediction remains a critical area for investigation.

**Thermodynamic features**, including siRNA intrinsic stability and siRNA-mRNA interaction energies (e.g., Gibbs free energy), have long been considered important (*8,15,16,21*). Tools like the ViennaRNA package (*22*) are frequently used for these calculations. However, the application of these features is not without pitfalls. For example, careful analysis suggests that some studies may have inadvertently used the incorrect siRNA strand (sense instead of antisense) for siRNA-mRNA Gibbs free energy calculations, leading to less accurate thermodynamic values (see Supplementary Fig. S1 for an example concerning La Rosa et al. (*15*)). Furthermore, dimensionality reduction techniques like Singular Value Decomposition (SVD) applied to thermodynamic profiles (*16*) might obscure interpretability or introduce dataset-specific artifacts.

Despite progress, systematically evaluating the contribution of different feature sets and complex architectures, especially with rigorous hyperparameter optimization and robust validation against issues like data leakage across gene targets, remains crucial. This study addresses this gap by systematically evaluating feature utility and architectural contributions. Through this process, I introduce RN.Ai-Predict, a model developed by first identifying the most predictive core features, primarily tokenized learned nucleotide embeddings, and then carefully assessing the additive value of more complex architectures. This bottom-up approach aims to build a powerful predictive model, emphasizing generalizability and robust performance.

## Dataset Compilation and Preprocessing

This study utilized a compiled siRNA dataset consisting of 3025 siRNA–mRNA pairs along with their corresponding experimentally determined efficacy values (Supplementary Fig S2). I sourced the compiled dataset from publicly available data used in La Rosa et al. (*15*) and Long et al. (*16*), which consolidated data from nine original studies: Harborth et al. (*23*), Huesken et al. (*13*), Katoh et al. (*24*), Khovorova et al. (*25*), Reynolds et al. (*1*), Sciabola et al. (*26*), Ui-Tei et al. (*2*), and Vickers et al. (*27*). In addition, data from Hsieh et al. (*3*) was extracted and tested.

For data partitioning, I split siRNA–mRNA pairs based on their associated target gene. I adopted this approach to prevent information leakage from siRNAs targeting the same gene (and thus potentially similar mRNA regions) between the training and test sets, thereby ensuring a more rigorous evaluation of model generalization. Gene-based partitioning also facilitated fairer comparisons with GNN models.

## Feature Engineering and Representation Evaluated

Based on the established importance of sequence and thermodynamic properties in siRNA efficacy, this study systematically evaluated the following feature categories and representations:

### 1. Nucleotide Sequence Representations

- *One-hot encoding*: I represented each nucleotide in the siRNA (19-21 nt) and its mRNA target context (19 nt flanking regions on both 5’ and 3’ ends of the target site) as a 4-dimensional binary vector, e.g. [0,1,0,0] for cytosine.
- *K-mer encoding*: I encoded siRNA sequences using 3-nucleotide k-mers, and mRNA context sequences using 4-nucleotide k-mers, following approaches similar to La Rosa et al. (*15*).
- *Pre-trained Tokenized Embedding (RNA-FM)*: I obtained embeddings for nucleotide sequences using the pre-trained RNA-FM model (*20*), as explored in OligoFormer (*17*).
- *Tokenized Learned Embedding*: I incorporated an embedding layer into the neural network to learn task-specific representations for each nucleotide. This layer generated a dense vector for every nucleotide in the siRNA and mRNA context sequences, optimizing the vectors’ positions in a multi-dimensional space based on the efficacy data. I treated the dimensionality of this vector space as a tunable hyperparameter.

For all sequence representations, the input consisted of the siRNA sequence and a 19-nucleotide flanking region of the mRNA on both the 5’ and 3’ sides of the siRNA binding site.

### 2. Thermodynamic Features

- *siRNA Thermodynamic Stability*: I calculated the internal thermodynamic stability for each siRNA duplex based on the nearest-neighbor parameters from Xia et al. (*21*).
- *siRNA-mRNA Interaction Gibbs Free Energy*: Using RNAup from the ViennaRNA package (*22*), I calculated the Gibbs free energy of binding between the antisense siRNA strand and its target mRNA, ensuring the correct strand was used for the interaction.

I evaluated these features both individually and in various combinations to determine their predictive power.

## Neural Network Architectures for Nucleotide Representation Processing

Beyond the choice of nucleotide representation, how the representation is processed within the neural network may impact efficacy prediction. I considered several common approaches for evaluation:

1. **Feed-forward Deep Neural Network (FFN)**: The model feeds the (flattened) nucleotide representation directly into a series of fully connected layers.
2. **Convolution Neural Networks (CNNs) to FFN**: CNNs apply convolution filters to extract local patterns. In the context of nucleotide sequences, 1D convolution can identify motifs or regional characteristics (e.g., GC-rich regions) by scanning along the sequence.
3. **Recurrent Neural Networks (RNNs) - LSTM/bi-LSTM to FFN**: Long Short-Term Memory (LSTM) networks, particularly bidirectional LSTMs (bi-LSTMs), are designed to capture sequential dependencies. Bi-LSTMs process the sequence in both forward and reverse directions, allowing the network to learn context from the entire sequence.
4. **Transformer Encoders to FFN**: Originating from NLP, transformer encoders use self-attention mechanisms to weigh the importance of different nucleotides in the sequence relative to each other, capturing long-range dependencies. This could be beneficial for identifying interactions between distant positions, such as the 5’ and 3’ ends of an siRNA or its target site.

I applied these architectures after representing nucleotides with a tokenized embedding layer.

## Systematic Evaluation of Features and Model Architectures

The predictive utility of features with potentially complex, non-linear relationships, such as nucleotide sequences, is often best assessed empirically. This study implemented a systematic approach to evaluate commonly used feature sets and neural network architectures for siRNA efficacy prediction. A key aspect of this approach was rigorous hyperparameter optimization for each feature-architecture combination to ensure fair comparisons.

Neural network models are highly sensitive to hyperparameter selection. Even simple architectures possess a vast hyperparameter space. For instance, a nine-layer feed-forward network, with four activation function choices per layer, 50 output size options, and three dropout levels, yields (4 × 50 × 3)^9^ possible configurations, an intractably large space for exhaustive search.

To effectively navigate this vast hyperparameter space, a two-stage hybrid optimization strategy was employed for each feature/architecture evaluation:

1. **Initial Broad Search with Hyperband**: I first utilized the Hyperband algorithm (*28, 29*) to efficiently explore a wide range of hyperparameter configurations. Hyperband works by adaptively allocating computational resources (e.g., training epochs) to different configurations, progressively promoting promising ones while quickly discarding those performing poorly. This initial stage served to broadly identify regions of the hyperparameter space likely to contain high-performing models. The specific ranges and step sizes for hyperparameters explored by Hyperband are detailed in Table 1 (“Hyperband Step/Choice” column).
2. **Refined Search with Bayesian Optimization**: Following the Hyperband search, I used the top ten performing hyperparameter configurations identified to initialize and constrain the search space for a subsequent Bayesian optimization phase. Bayesian optimization builds a probabilistic surrogate model of the objective function (e.g., validation R^2^ or MSE) and uses an acquisition function to intelligently select the next set of hyperparameters to evaluate, balancing exploration of uncertain regions with exploitation of known good regions (*30*). This allows for a more fine-grained search within the promising areas identified by Hyperband. For example, while Hyperband might explore dense layer output sizes in coarse steps (e.g., 100, 200, 300), Bayesian optimization could refine this to single-unit precision (see Table 1, “Bayesian Step/Float/Choice” column).

**Table 1:** Hyperparameter search space for optimization. Each row details a hyperparameter, the range or choices explored, and the step size or method for Hyperband and Bayesian optimization stages.

| Hyperparameter | Choices/Range | Hyperband Step | Bayesian Step |
| --- | --- | --- | --- |
| <u>FFN Hyperparameters</u> |  |  |  |
| DNN layers | 1-9 | 1 | 1 |
| Activation | relu, tanh, elu, linear, mish | Choice | Choice |
| Output units | 100-500 | 100 | 1 |
| Dropout | 0.0, 0.1, 0.2, 0.3 | Choice | Float |
| <u>Nucleotide Representation Hyperparameters</u> |  |  |  |
| Embedding output | 1, 2, 3, 4, 8, 16, 32, 64, 128, 256 | Choice | 1 |
| Dropout | 0.0, 0.1, 0.2, 0.3 | Choice | Float |
| <u>Small FFN (Combination Post Architecture)</u> |  |  |  |
| DNN layers | 0-3 | 1 | 1 |
| Dense layer output | 100-500 | 100 | 1 |
| Dropout | 0.0, 0.1, 0.2, 0.3 | Choice | Float |
| <u>Convolution 1D Hyperparameters</u> |  |  |  |
| Filters | 1, 2, 3, 4, 8, 16, 32, 64, 128, 256 | Choice | 1 |
| Kernel Size | 2-16 | 1 | 1 |
| Activation | relu, tanh, elu, linear, mish | Choice | Choice |
| <u>Bi-LSTM Hyperparameters</u> |  |  |  |
| LSTM output units | 5-500 | 25 | 1 |
| Activation | relu, tanh, elu, linear, mish | Choice | Choice |
| <u>Transformer Encoder Hyperparameters</u> |  |  |  |
| Encoders | 1-5 | 1 | 1 |
| Attention heads | 1, 2, 4, 8 | Choice | 1 |
| Encoder dropout | 0.0, 0.1, 0.2, 0.3 | Choice | Float |
| FFN dropout | 0.0, 0.1, 0.2, 0.3 | Choice | Float |
| Flatten choice | Flatten, Average, Max | Choice | Choice |

For the Bayesian optimization phase, a minimum of *N*_*HP*_ initial random points were sampled to seed the surrogate model, where *N*_*HP*_ is the number of hyperparameters being tuned (typically set to 4 × *N*_*HP*_ in this study). The optimization then proceeded for a number of trials equal to the maximum of 500 or 12 × *N*_*HP*_.

During the training of each model configuration by Bayesian optimization, I employed a learning rate scheduling strategy. The initial learning rate was reduced by a factor of ten if the validation set Mean Squared Error (MSE) did not improve for 10 consecutive epochs. I halted training (early stopping) if the validation MSE showed no improvement for 20 consecutive epochs, to prevent over-fitting and save computational resources.

This hyperparameter optimization process itself is sensitive to the choice of validation data. To ensure robustness, I generated five distinct sets of optimized hyperparameters for each feature/architecture combination under evaluation. I achieved this by performing the two-stage (Hyperband + Bayesian) optimization five times, each time using a different fold of a 5-fold gene-based cross-validation split of the (training) data as the validation set for that optimization run (Fig. 1A). This resulted in five distinct optimized hyperparameter profiles.

**Figure 1:**
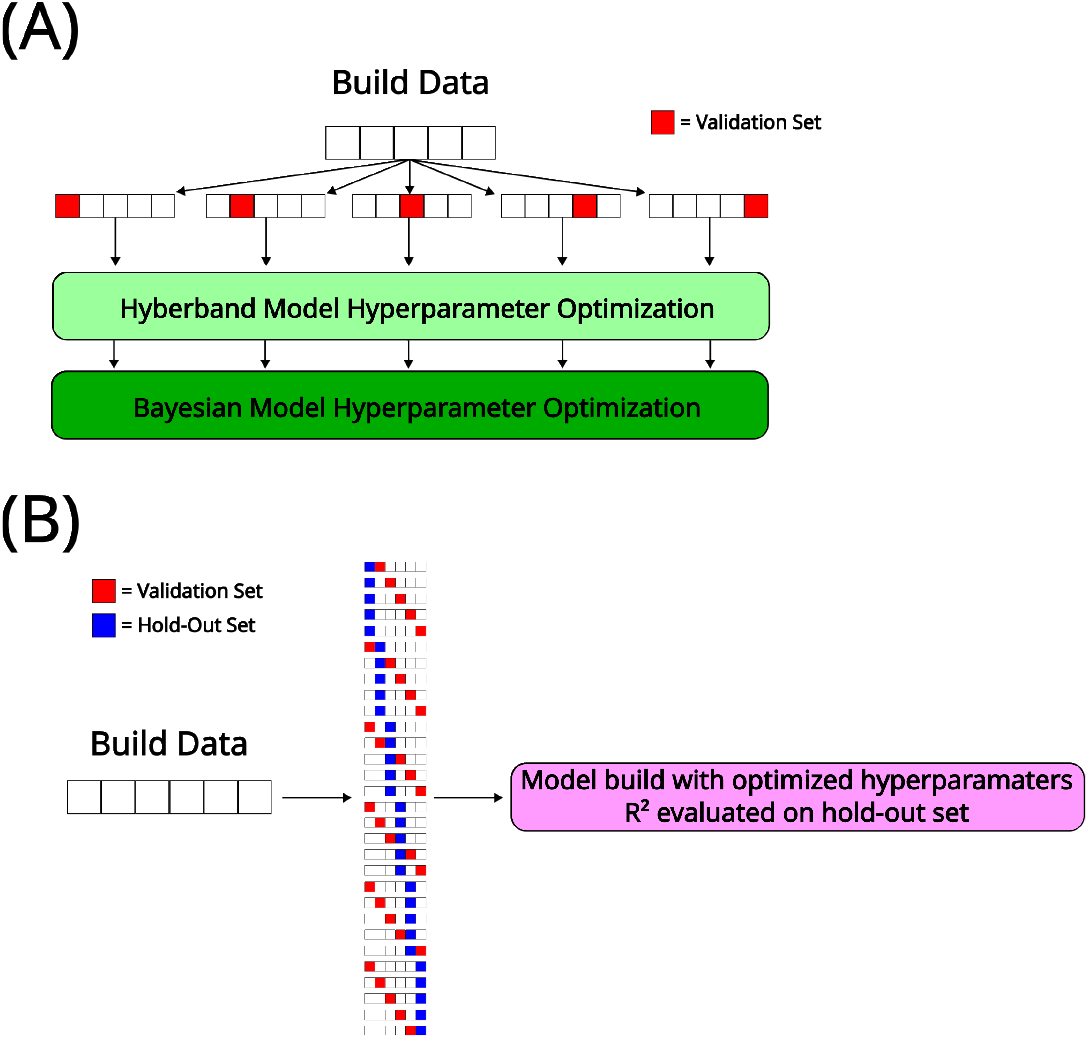
Strategy for systematic feature and architecture evaluation. An evaluation framework was used to assess different features and AI architectures. (A) Hyperparameter optimization schematic: For each feature/architecture, five distinct hyperparameter optimization processes were run. Each process used a different fold of a 5-fold split of the training data as its validation set for tuning (initially with Hyperband, then refined with Bayesian optimization). The top Bayesian optimized hyperparameters were selected from each of these five processes. (B) Performance assessment schematic: Each selected hyperparameter configuration was then evaluated using a 6-fold gene-based cross-validation. In each fold, the model was trained (on 4 gene-folds), validated for early stopping (on 1 gene-fold), and its R^2^ score computed on a held-out gene-fold. This yields 30 R^2^ values per hyperparameter set, resulting in many R^2^ values (5×30 = 150) for each feature/architecture, ensuring assessment of generalizability to unseen genes and robustness of hyperparameters.

I then assessed each of these candidate hyperparameter sets for generalizability using a separate 6-fold gene-based cross-validation scheme (Fig. 1B). In each iteration of this 6-fold CV, four folds were used for training, one fold for validation (primarily for early stopping at convergence, as hyperparameters were fixed), and the remaining fold was held out as a test set for final performance assessment. This extensive process, repeated across the 5 optimized hyperparameter sets and 6 CV folds, generates 150 R^2^ values (5 × 6 × 5 = 150) per model configuration, providing a robust statistical distribution of performance and guarding against spurious results from a single train/test split.

I primarily evaluated model performance using the coefficient of determination (R^2^), along with Pearson correlation coefficient (PCC), mean absolute error (MAE), and mean squared error (MSE). The distribution of R^2^ values obtained from the hold-out sets was analyzed to identify the most predictive feature sets and model architectures for constructing the final RN.Ai-Predict model. A complete list of hyperparameters explored is provided in Table 1.

## Results: Data Source Evaluation and Normalization

The Huesken et al. dataset (*13*), a large component of the compiled data, contained knockdown efficiencies exceeding 100% (Fig. 2A), which are biologically implausible. I tested two normalization strategies for the Huesken efficacy values: capping at 100% (efficacy=1.0), and min-max scaling (*17*) (Fig. 2B). I trained models using tokenized learned embeddings (Fig. 2C), with hyperparameters optimized via 5-fold CV (Fig. 2D), on these normalized Huesken datasets and tested the models on other public datasets.

**Figure 2:**
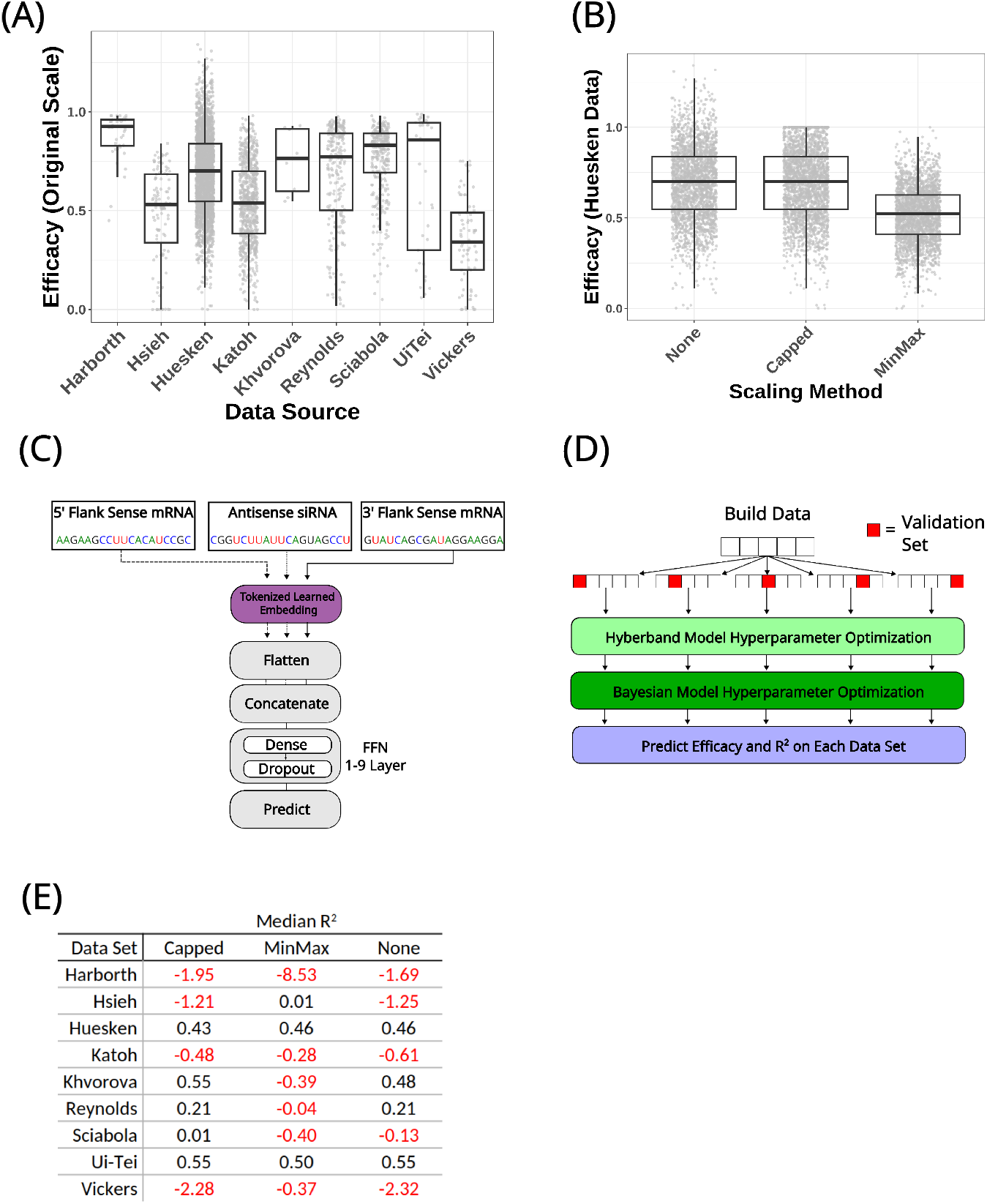
Dataset evaluation and impact of normalization. (A) Distribution of siRNA efficacy values (knockdown proportion) across individual public datasets. (B) Distribution of efficacy values from the Huesken dataset: original, after Min/Max scaling to [0,1], and after capping values at 1.0 (100% knockdown). (C) Overview of the tokenized learned embedding nucleotide model (siRNA sequence and 19nt mRNA flanks fed to an FFN) used to evaluate dataset performance and normalization strategies. (D) Schematic of the hyperparameter tuning process used to evaluate R^2^ values for each dataset. (E) Median R^2^ values obtained when models trained on differently normalized versions of the Huesken dataset were tested on all public datasets.

Capping efficacy values at 1.0 for the Huesken data yielded more consistent and robust crossdataset generalization (Fig. 2E). Models trained on capped Huesken data produced positive median R^2^ values on 5 out of 9 other datasets, compared to 3 out of 9 using min-max normalization. This strategy also preserved performance on the Reynolds et al. dataset (*1*). The Harborth, Hsieh, Katoh, and Vickers datasets showed poor concordance (*R*^2^ *<* 0) with models trained on Huesken data, suggesting significant divergence. I excluded these four datasets from subsequent primary modeling efforts to avoid bias, though their impact was later re-evaluated.

## Results: Identifying the Core Predictive Feature: Nucleotide Representation

I evaluated four nucleotide representation approaches (One-hot, k-mer, pre-trained RNA-FM, and tokenized learned embedding) using a baseline FFN architecture (Fig. 3A), with hyperparameters optimized and performance assessed as per Fig. 1.

**Figure 3:**
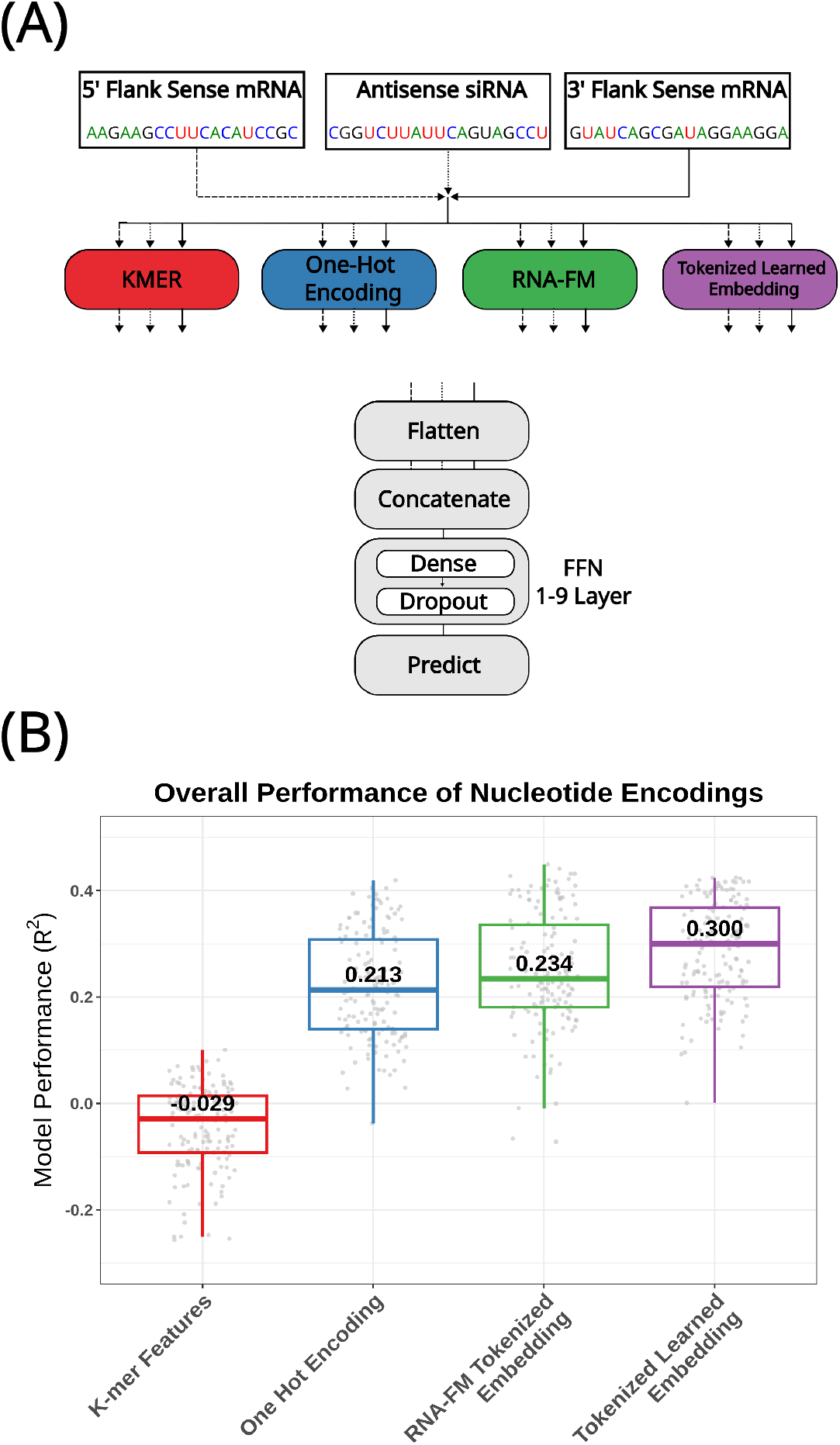
Evaluation of nucleotide features. Hyperparameters for all models were optimized and evaluated using the strategy in Fig. 1. (A) Schematic representation used for evaluating different nucleotide representations (K-mer, One-hot, RNA-FM, Tokenized Learned Embedding). (B) Box plots of R^2^ values for each nucleotide encoding strategy. Median R^2^ values are indicated within the plot area.

The evaluation of nucleotide representations revealed a clear performance hierarchy (Fig. 3B). Tokenized learned embedding processed by a simple feed-forward network (FFN) achieved the highest predictive performance, with a median R^2^ of 0.300. This outperformed pre-trained RNA-FM embeddings (R^2^=0.234), standard one-hot encoding (R^2^=0.213), and k-mer encoding, which failed to generalize (R^2^=-0.029). This result underscores that both positional context (lost in k-mers) and the ability to learn task-specific nucleotide relationships (absent in one-hot, potentially suboptimal in pre-trained RNA-FM) are crucial for efficacy prediction.

To further validate the power of the embedding layer itself, I tested whether a one-hot encoded input could match its performance if made functionally equivalent. Notably, recent deep learning models such as GNN4siRNA, siRNADiscovery, and OligoFormer typically feed one-hot encodings directly into non-dense layers like GNNs or convolutions (*15–17*). In contrast, I explored adding an initial dense layer directly to the one-hot input, effectively creating a learnable lookup table. This simple architectural change improved performance to a median R^2^ of 0.280 (Supplementary Fig. SS4). However, this still did not surpass the dedicated token-based embedding, confirming it as the most effective approach.

Due to the superior performance of tokenized embeddings fed to a simple FFN, I selected this architecture as my final model, which I term RN.Ai-Predict. All subsequent analyses use tokenized embedding for nucleotide representation.

## Results: The Additive Value of Thermodynamic and Architectural Complexity

Having established the baseline performance of RN.Ai-Predict using only sequence embeddings, I next investigated whether incorporating thermodynamic features or more complex architecture could provide additional predictive power.

First, I evaluated siRNA stability and siRNA-mRNA Gibbs free energy, both alone and in combination with the learned embeddings (Fig. 4A). While siRNA stability showed some predictive capability on its own (median R^2^ = 0.191), adding any thermodynamic features to the RN.Ai-Predict architecture did not improve performance beyond the 0.300 median R^2^ achieved with embeddings alone (Fig. 4B). This suggests the learned embeddings implicitly capture the relevant predictive information contained in these properties.

**Figure 4:**
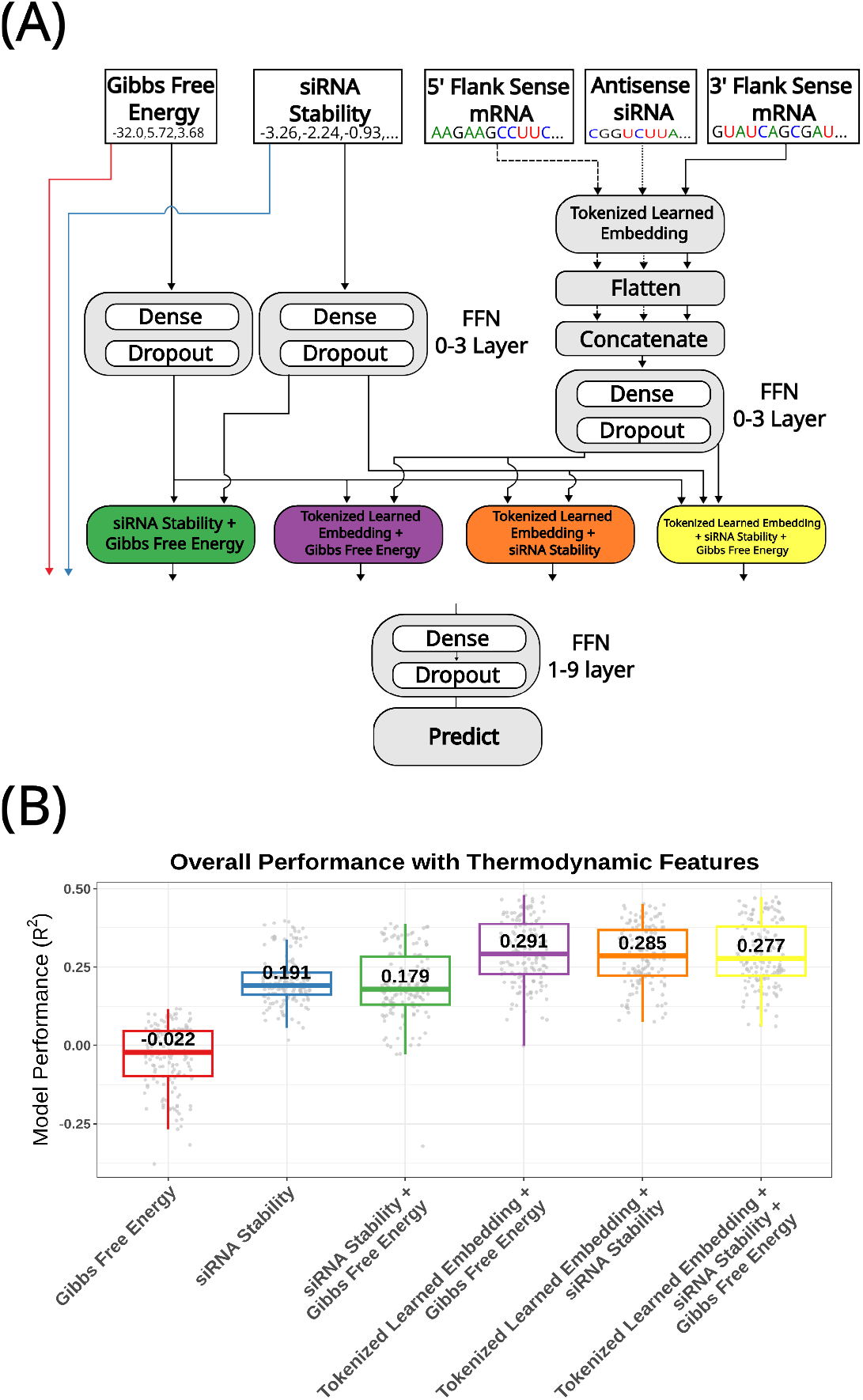
Evaluation of thermodynamic features. Hyperparameters for all models were optimized and evaluated using the strategy in Fig. 1. (A) Schematic representation of models used to evaluate thermodynamic features (siRNA stability, siRNA-mRNA Gibbs free energy) alone or in combination with tokenized learned embeddings. (B) Box plots of R^2^ values for models using thermodynamic features.

Next, I tested whether more complex neural network architectures could extract additional information from the learned embeddings. Two main architectural configurations were tested for each of these advanced layers (Fig. 5A illustrates the general scheme):

1. **Sequential Processing**: I fed the tokenized learned embeddings of the siRNA and mRNA sequence (target site plus flanks) directly into the Conv1D, bi-LSTM, or Transformer Encoder layers. Including the target site within the mRNA, not just the flanking regions, was a deliberate choice to allow these advanced architectures to identify any predictive signals across the binding region and through the flanks. The output of this specialized layer was then passed to dense layers for efficacy prediction.
2. **Parallel Processing with Concatenation**: To ensure that local features captured by a direct FFN on embeddings were not lost, I also tested a parallel architecture. Here, the output from the specialized layer (Conv1D, bi-LSTM, or Transformer processing the embeddings) was concatenated with a separate, independently processed tokenized learned embedding of the siRNA and flanking mRNA. This combined feature set was then fed to final dense layers for prediction. This approach aimed to see if the specialized architectures could learn complementary information (e.g., broader sequential patterns or distal dependencies) to the highly effective local features learned directly by the embedding-to-FFN pathway.

**Figure 5:**
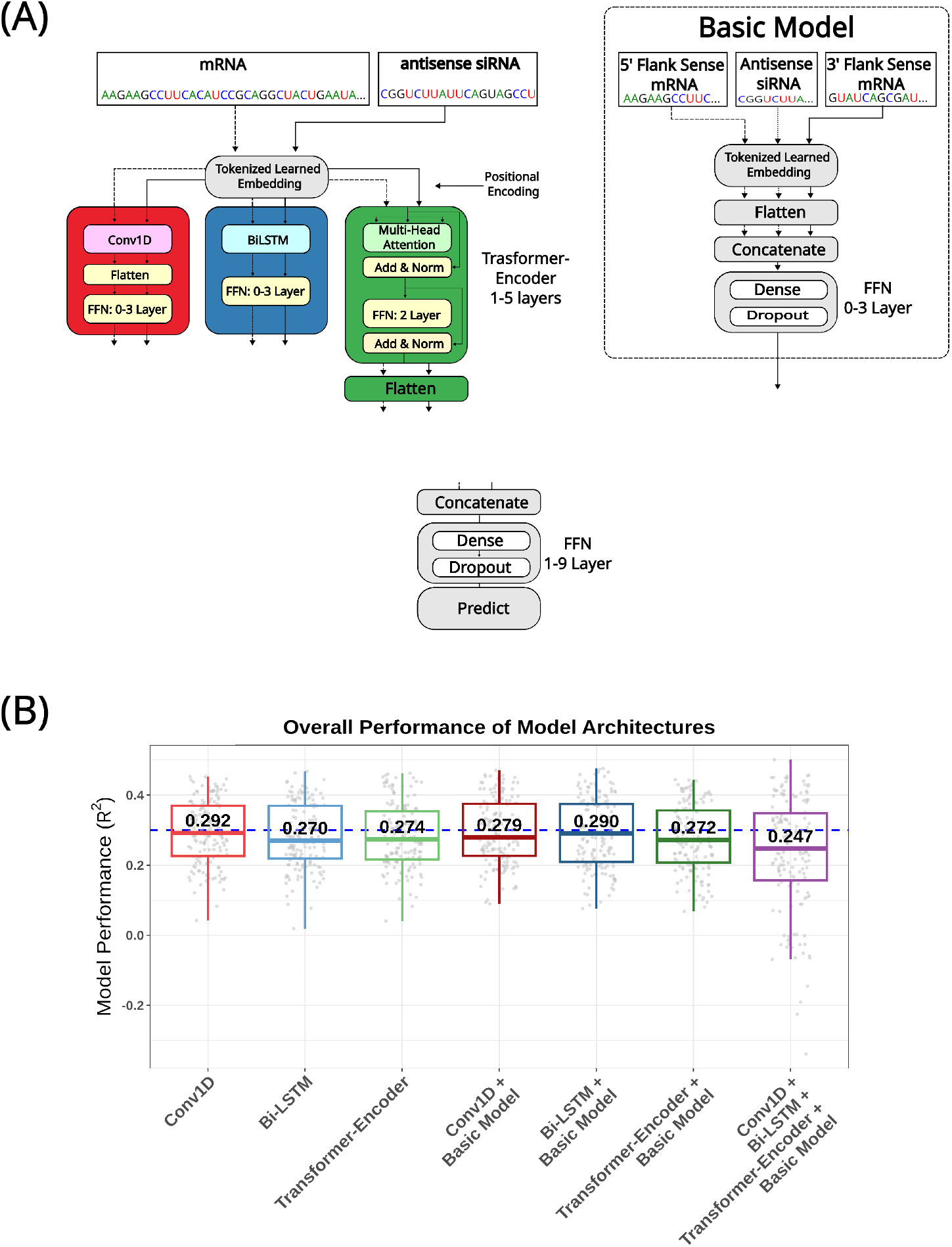
Evaluation of advanced neural network architectures for sequence processing. Hyperparameters for all models were optimized and evaluated using the strategy in Fig. 1. (A) Schematic of architectures tested. (B) Box plots of R^2^ values for models incorporating Conv1D, bi-LSTM, or Transformer Encoder layers on top of tokenized learned embeddings. The dashed line indicates the median R^2^ (0.300) achieved by tokenized learned embeddings alone (from Fig. 3B).

When used in a **sequential processing** manner, where the specialized architecture directly processed the embeddings, the median R^2^ values were:

- Tokenized embedding → Conv1D layers → FFN: 0.292
- Tokenized embedding → bi-LSTM layers → FFN: 0.270
- Tokenized embedding → Transformer Encoder layers → FFN: 0.274 None of these configurations outperformed RN.Ai-Predict (Fig. 5B).

Next, I evaluated the **parallel processing with concatenation** approach. While this strategy slightly improved the performance of some specialized architectures compared to their sequential-only counterparts, none of the combined models meaningfully surpassed the predictive power of RN.Ai-Predict:

- (Tokenized embedding → Conv1D) + (Tokenized embedding) → FFN: 0.279
- (Tokenized embedding → bi-LSTM) + (Tokenized embedding) → FFN: 0.290
- (Tokenized embedding → Transformer Encoder) + (Tokenized embedding) → FFN: 0.272 In addition to these, I tested a model with all 3 architectures (Conv1D, Bi-LSTM, and Transformer-Encoder) concatenated with an independently processed tokenized learned embedding, which achieved a median R^2^ of 0.247 (Fig. 5B). Overall, no models with specialized architecture outperformed RN.Ai-Predict, indicating no substantial gain from the added complexity.

## Results: Benchmarking RN.Ai-Predict Against Other Models

### Comparison with Legacy Models

Since the end-to-end learned tokenized embedding of RN.Ai-Predict gave the best results, I tested its performance against other siRNA prediction methods: first-generation rules, second-generation ML models, and current deep learning models. To facilitate this comparison, I used the optimized hyperparameters derived from my first validation fold (Supplementary Tab. S1) to train new models, each holding out one of the specific genes as a test set. These genes were selected based on their data coverage and their exclusion from the training data of the legacy models. For the third-generation models, their publicly available code allowed applying my exact hold-out strategy, ensuring a direct comparison on unseen data.

I benchmarked RN.Ai-Predict against two widely recognized legacy models: the Reynolds ruleset and the DSIR algorithm (http://biodev.cea.fr/DSIR/)(*1, 7*). For DSIR, I evaluated both its standard ‘Score’ and its ‘Corrected score’ outputs. The comparison was performed on a set of five hold-out genes that were not present in the original training or development sets of these legacy models. Performance was measured by the Spearman correlation between each model’s predicted score and the experimentally measured siRNA efficacy. For RN.Ai-Predict, I report the performance distribution of the five models generated by 5-fold CV, each using a different validation fold for creation.

The results, summarized in Table 2 and visualized in Figure 6A, demonstrate that RN.Ai-Predict consistently outperforms both legacy models on these unseen gene targets. The legacy models showed inconsistent and often poor performance. For instance, while the Reynolds Score was comparable to RN.Ai-Predict on a single gene (HK2, with a Spearman correlation of 0.27 vs. 0.25), it struggled elsewhere. The DSIR algorithm failed to generalize almost entirely, yielding negative correlations for most genes. In stark contrast, RN.Ai-Predict achieved a positive correlation for every gene tested, with over half showing a strong median correlation above 0.50. This performance gap highlights the superior generalizability of the learned embedding approach compared to models based on fixed rules.

**Table 2:** Comparative performance of RN.Ai-Predict and legacy models on seven hold-out genes. Performance is measured by the Spearman Correlation Coefficient. For RN.Ai-Predict, the median and range [min–max] of the five cross-validation models are shown. The best-performing model for each gene is highlighted in bold.

| Hold-out Gene | RN.Ai-Predict | Reynolds | DSIR | DSIR |
| --- | --- | --- | --- | --- |
|  | Median [Min–Max] | Score | (Score) | (Corrected) |
| TMEM63B | **0.60 [0.26 – 0.62]** | 0.24 | N/A | N/A |
| DENND5A | **0.53 [0.30 – 0.58]** | 0.13 | N/A | N/A |
| HPSE | **0.22 [0.10 – 0.26]** | -0.11 | -0.23 | -0.28 |
| HIF1A | **0.42 [0.16 – 0.45]** | 0.15 | 0.00 | -0.13 |
| HK2 | 0.25 [0.07 – 0.27] | 0.27 | **0.33** | -0.08 |
| hCyPB | **0.52 [0.13 – 0.57]** | N/A | -0.07 | -0.07 |
| GAPDH | **0.61 [0.46 – 0.71]** | N/A | -0.02 | 0.01 |

**Figure 6:**
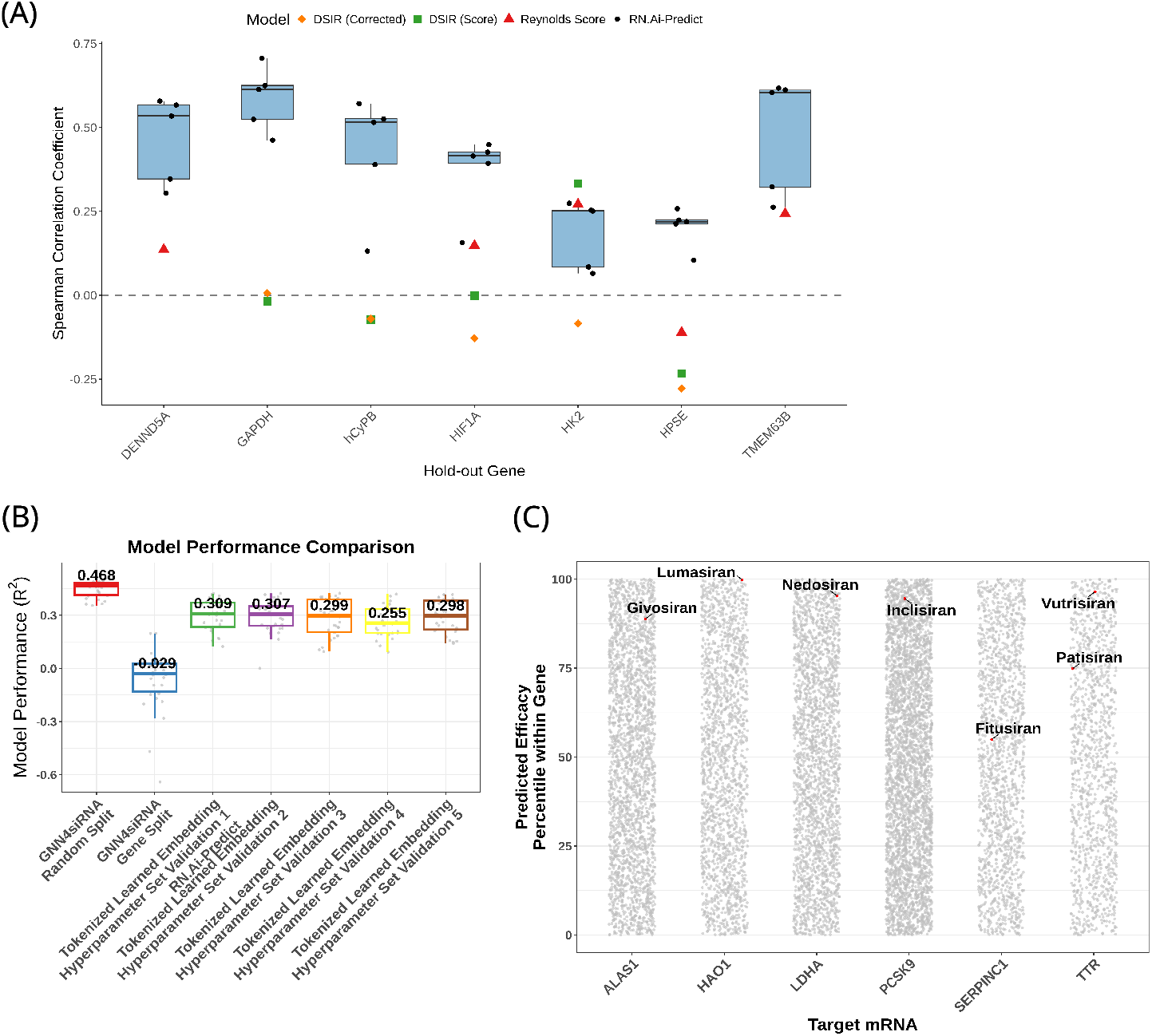
Comparative performance of the optimized Tokenized Learned Embedding model (RN.Ai-Predict). (A) Spearman Correlation Coefficient values for RN.Ai-predict’s, Reynolds Score, and DSIR original and corrected score. For RN.Ai-Predict, training was done on a dataset where the indicated gene was held-out then tested. (B) R^2^ values for tokenized learned embedding (separated by hyperparameter set derived from the 5 different validation sets, with the best performing labeled as RN.Ai-predict) compared to GNN4siRNA. Both were evaluated using 6-fold genebased cross-validation. GNN4siRNA was also tested with random siRNA splits (“GNN4siRNA Random Split”) as per its original publication. (C) Application of RN.Ai-Predict to predict efficacy for all possible siRNAs targeting mRNAs of five FDA-approved therapeutics. The actual therapeutic siRNA sequences (Givosiran, Lumasiran, Nedosiran, Inclisiran, Fitusiran, Vutrisiran, and Patisiran) are shown by red dots, plotted by their predicted efficacy percentile within their respective gene.

### Comparison with GNN-based Models

To further benchmark RN.Ai-Predict, a comparison was made against GNN4siRNA, a recent graph neural network model (*15*). I selected GNN4siRNA due to its linear prediction of efficacy and the public availability of its code, the latter allowing for re-evaluation on strictly unseen gene sets. I evaluated GNN4siRNA code (available at https://github.com/Roco-scientist/ GNN4siRNA-predict with minor modifications for this comparative analysis) under two data splitting schemes:

1. **Random siRNA split**: siRNAs randomly assigned to training/test sets, as per the original GNN4siRNA publication.
2. **Gene-based split**: siRNAs split by gene, ensuring all siRNAs for a given gene are in the same set (train or test), consistent with the evaluation methodology in this study to prevent information leakage.

As previously noted (Supplementary Fig. SS1 and “Feature Engineering and Representation” section), the original GNN4siRNA study appeared to use incorrect thermodynamic calculations. This was corrected to create a better comparison.

When GNN4siRNA was evaluated with random siRNA splitting (Scheme 1), it achieved a median R^2^ of 0.468 on the hold-out sets, consistent with the performance reported by its original authors (Fig. 6B). However, when evaluated using the more stringent gene-based split, GNN4siRNA’s median R^2^ reduced to -0.029, indicating it failed to generalize to unseen genes and performed equivalent to a random predictor.

In contrast, the RN.Ai-Predict model (derived from validation set 1) achieved a median R^2^ of 0.309 on the same gene-based split (Fig. 6A). This highlights RN.Ai-Predict’s superior generalizability to new gene targets. The figure also shows the performance distribution for RN.Ai-Predict models derived from each of the five distinct hyperparameter optimization runs, demonstrating consistent positive predictive performance across them.

### Application to FDA-Approved siRNA Therapeutics

The GNN4siRNA results, particularly its high performance only when genes are seen during training, suggest it might be leveraging gene-specific information. To explore if a gene-relative ranking could be useful, I used the RN.Ai-Predict model to predict efficacy for all possible siRNAs sequences (matching the length of the therapeutic) targeting the mRNAs of seven FDA-approved siRNA therapies: Givosiran, Lumasiran, Nedosiran, Inclisiran, Fitusiran, Vutrisiran, Patisiran. The predicted efficacy values for siRNAs within each gene were then converted to percentiles.

Remarkably, the actual sequences of Givosiran, Lumasiran, Nedosiran, Inclisiran, Vutrisiran, Patisiran all scored above the 75th percentile within their respective target genes (Fig. 6C). Fitusiran was the only therapy to fall below the 75th percentile. This is notable as RN.Ai-Predict was not trained on data including chemically modified siRNAs, yet it successfully ranked these highly effective therapeutic sequences favorably based solely on their nucleotide composition and mRNA context.

## Discussion

This study demonstrates that for siRNA efficacy prediction, a systematically optimized model based on learned nucleotide embeddings can achieve greater generalizability than more complex architectures. I developed my resulting model, RN.Ai-Predict, through a rigorous evaluation of feature representations and architectural complexity, placing it in direct comparison with the current state-of-the-art models. While previous models have made significant contributions, this work provides

a detailed comparative analysis against contemporary approaches, particularly emphasizing performance within a stringent, gene-based evaluation framework designed to prevent data leakage. The current landscape of advanced models includes GNN-based approaches like GNN4siRNA and siRNADiscovery (*15, 16*), and transformer-based models like OligoFormer, which incorporates pre-trained RNA-FM embeddings (*17*). My analysis shows that the simpler architecture of RN.Ai-Predict outperforms GNNs, transformer encoders, and models using pre-trained RNA-FM embeddings on the critical task of generalizing to unseen gene targets.

The application of GNNs to RNAi efficacy prediction, while innovative, may face inherent limitations. GNNs typically model siRNAs and mRNA targets as nodes with edges representing interactions. This relies on comprehensive interaction data, but in practice, complete off-target profiles for all siRNAs are rarely known. Thus, the constructed graph often represents only a small, potentially biased, subset of the true biological interaction network. My findings support this concern: GNN4siRNA performed well when siRNAs targeting genes seen in training were included in the test set (via random siRNA splitting) but failed to generalize to entirely new genes, an issue only revealed by the strict gene-based cross-validation strategy in this study. This suggests GNNs might capture gene-specific patterns (perhaps related to mRNA abundance or structure, for which expression level data was largely unavailable in the compiled dataset) rather than universally generalizable sequence features of siRNAs predictive of efficacy on novel targets.

Regarding other advanced neural network architectures, this study found that applying Conv1D, bi-LSTM, or Transformer Encoder layers on top of tokenized learned embeddings did not yield superior performance compared to a simpler FFN alone. This contrasts with some complex models like OligoFormer, which sequentially stack several such architectures (*17*). My results suggest that extensive downsampling or complex sequential processing may not be beneficial and could even be detrimental if the initial embedding already captures sufficient information, especially with datasets of the current scale. Overspecification of model architecture can lead to higher variance and poorer generalization if not supported by commensurate increases in data or signal.

A key finding of this study is the superior performance of end-to-end tokenized learned embeddings compared to pre-trained embeddings (RNA-FM) and other traditional representations like one-hot or k-mer encoding for this specific task. While OligoFormer utilizes RNA-FM (*17*), my results (Fig. 3B) showed that embeddings learned directly for siRNA efficacy prediction (median R^2^=0.300) outperformed RNA-FM (median R^2^=0.234). This suggests that task-specific embeddings, optimized from the data, can better capture the nuanced sequence determinants of siRNA activity.

The RN.Ai-Predict model, based on tokenized learned embeddings and an optimized FFN, demonstrated robust performance and generalizability to unseen genes. Its ability to rank FDA-approved siRNA sequences highly, despite not accounting for chemical modifications, further underscores the power of learning relevant sequence features from the available data.

## Limitations and Future Directions

This study, while comprehensive, has limitations. The dataset, though compiled from multiple sources, is still of moderate size for deep learning and predominantly features unmodified siRNAs. The exclusion of certain datasets due to inconsistency, while necessary for robust modeling, means the model is not trained on the full spectrum of available data. Future work should focus on:

1. **Incorporating Chemical Modifications**: Extending the model to account for common siRNA chemical modifications is crucial for predicting the efficacy of therapeutically relevant molecules.
2. **Larger and More Diverse Datasets**: Expanding the training dataset with more high-quality, consistently processed data, including diverse siRNA designs and target genes, will be essential.
3. **Exploring Gene-Specific Effects**: Investigating the gene-specific signals that GNNs appear to capture (e.g. expression levels if such data becomes available alongside efficacy) could lead to hybrid models that combine the strengths of sequence-based prediction with gene-context information.
4. **Prospective Validation**: The ultimate test for any predictive model is its ability to guide the design of novel, highly effective siRNAs, which requires prospective experimental validation.

In conclusion, this systematic evaluation demonstrates that a well-optimized neural network utilizing tokenized learned embeddings for sequence representation can achieve strong predictive performance for siRNA efficacy, offering a robust and generalizable foundation for computational siRNA design. This approach prioritizes effective feature learning and rigorous model validation over sheer architectural complexity.

## Author contributions

RTC did all work within this paper.

## Data and materials availability

Model and code is available at https://github.com/Roco-scientist/RN.Ai-Predict.

## Supplementary materials

Figs. S1 to S8

Tables S1

