## Supplemental Data for "Systematic feature and architecture evaluation reveals tokenized learned embeddings enhance siRNA efficacy prediction"

Rory Coffey*

**This PDF file includes:**

Figures S1 to S8

Tables S1

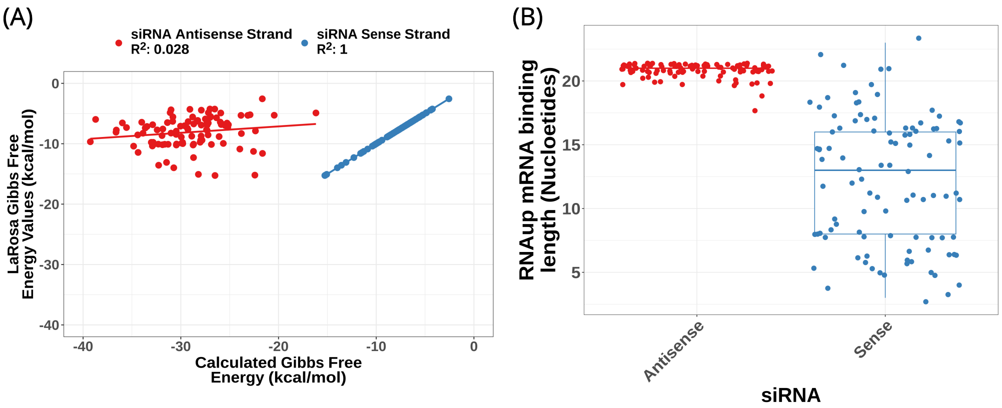


**Figure S1**: **Gibbs Free Energy of siRNA-mRNA Interaction: La Rosa et al. Used the siRNA Sense Strand Instead of the Antisense Strand.** (A) Gibbs free energy was calculated for 100 sample siRNA-mRNA pairs from the La Rosa et al. GNN study. Calculations were performed for both the siRNA antisense strand (red) and the sense strand (blue), with the calculated values plotted on the x-axis and La Rosa's reported values on the y-axis. The results demonstrate a perfect correlation ($R^2$ = 1) between La Rosa's calculations and the sense strand values, confirming that the sense strand was used. In contrast, the antisense strand calculations show no significant correlation ($R^2$ = 0.028) with La Rosa's values. Moreover, the calculated antisense strand exhibits more negative Gibbs free energy values, indicating stronger and more favorable siRNA-mRNA interactions, which is expected when the correct strand is used for binding. (B) The length of the mRNA bound to siRNA reported by RNAup for both the sense and antisense siRNA strand. Median value for antisense strand indicates 21 nucleotides, the expected value for a perfect match.

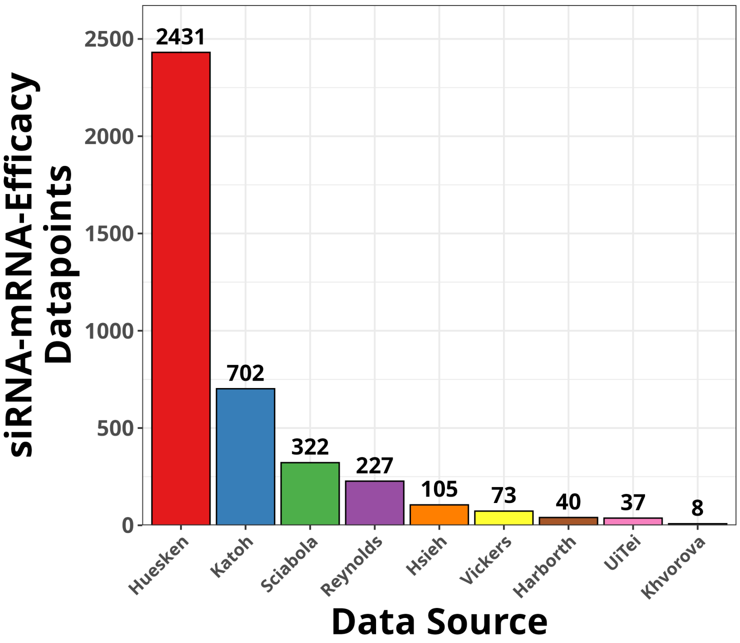


**Figure S2**: **Number of siRNA-mRNA-Efficacy Data points Per source Dataset.** Total number of datapoints per dataset. The final used datasets were Huesken (2431), Khvorova (8), Reynolds (227), Scaibola (322), and UiTei (37) for a total of 3025 of the 3945 available datapoints.

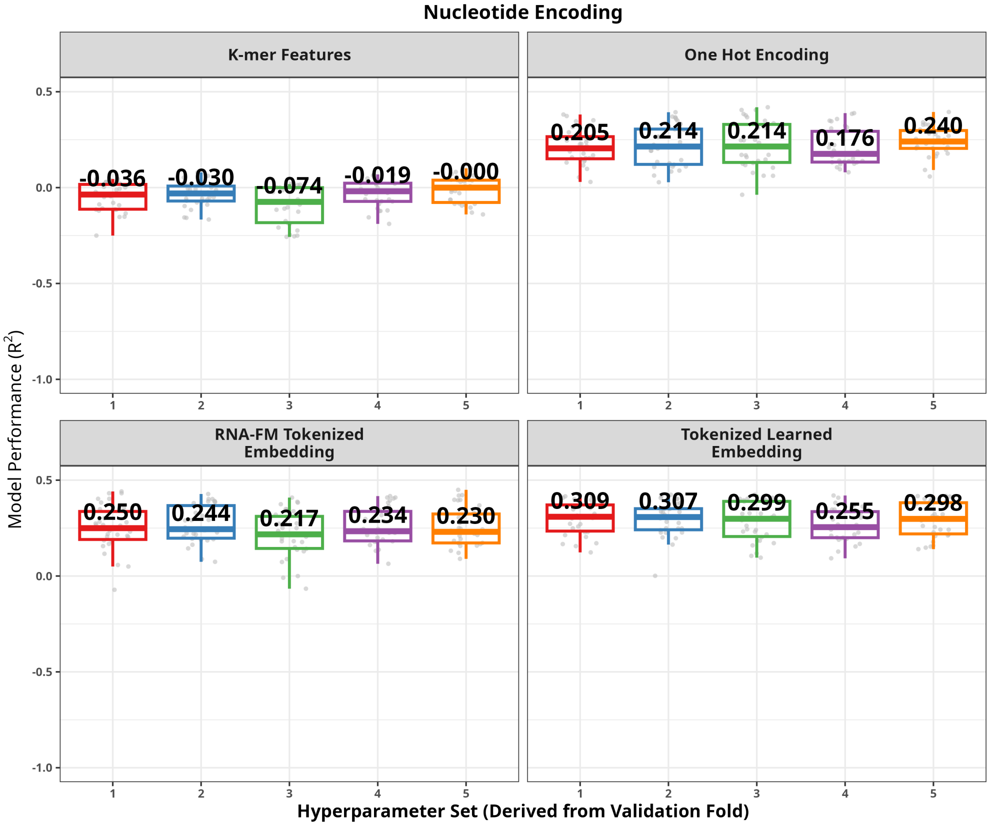


**Figure S3**: **Nucleotide Feature Evaluation Split by Validation Set.** With the hyperparameter optimization scheme from Fig 1, 5 different validation datasets produced 5 different hyperparameter sets. Each graph has the median $R^2$ on hold-out data labeled. Each evaluated hyperparameter set aggregated in Fig 3B.

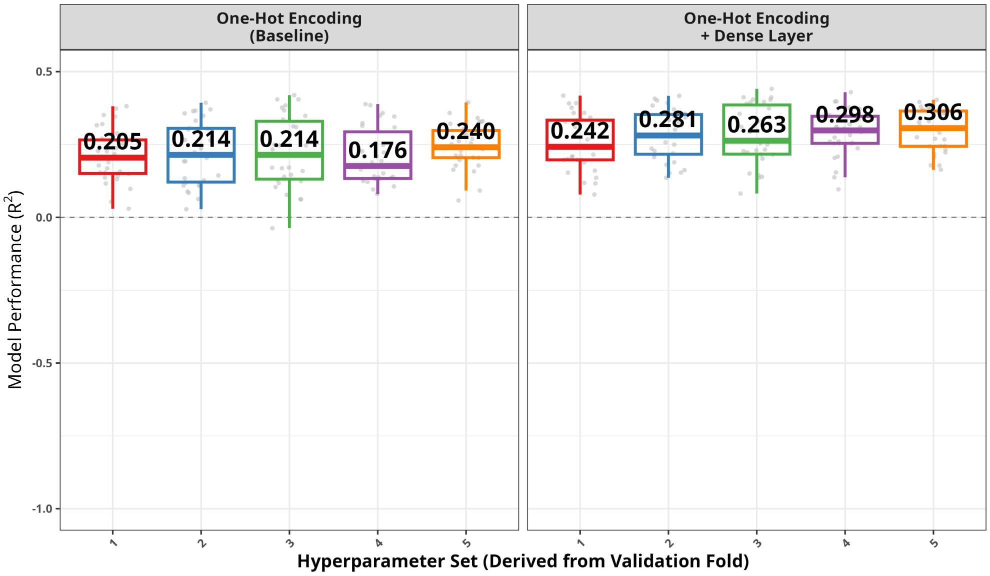


**Figure S4**: **Adding a Dense Layer After One-Hot Encoding Enhances Model Performance.** With the hyperparameter optimization scheme from Fig 1, 5 different validation datasets produced 5 different hyperparameter sets. Each graph has the median $R^2$ on hold-out data labeled. The overall median $R^2$ of One-Hot Encoding Baseline is 0.213 and with the added dense layer is 0.280

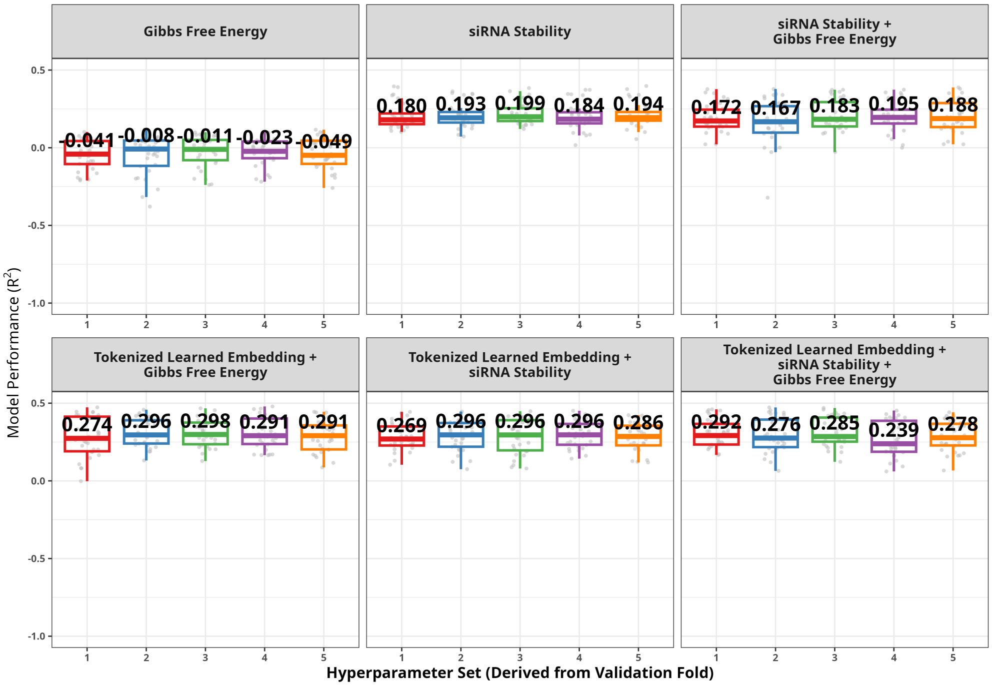


**Figure S5**: **Thermodynamic Feature Evaluation Split by Validation Set.** With the hyperparameter optimization scheme from Fig 1, 5 different validation datasets produced 5 different hyperparameter sets. Each graph has the median $R^2$ on hold-out data labeled. Each evaluated hyperparameter set aggregated in Fig 4B.

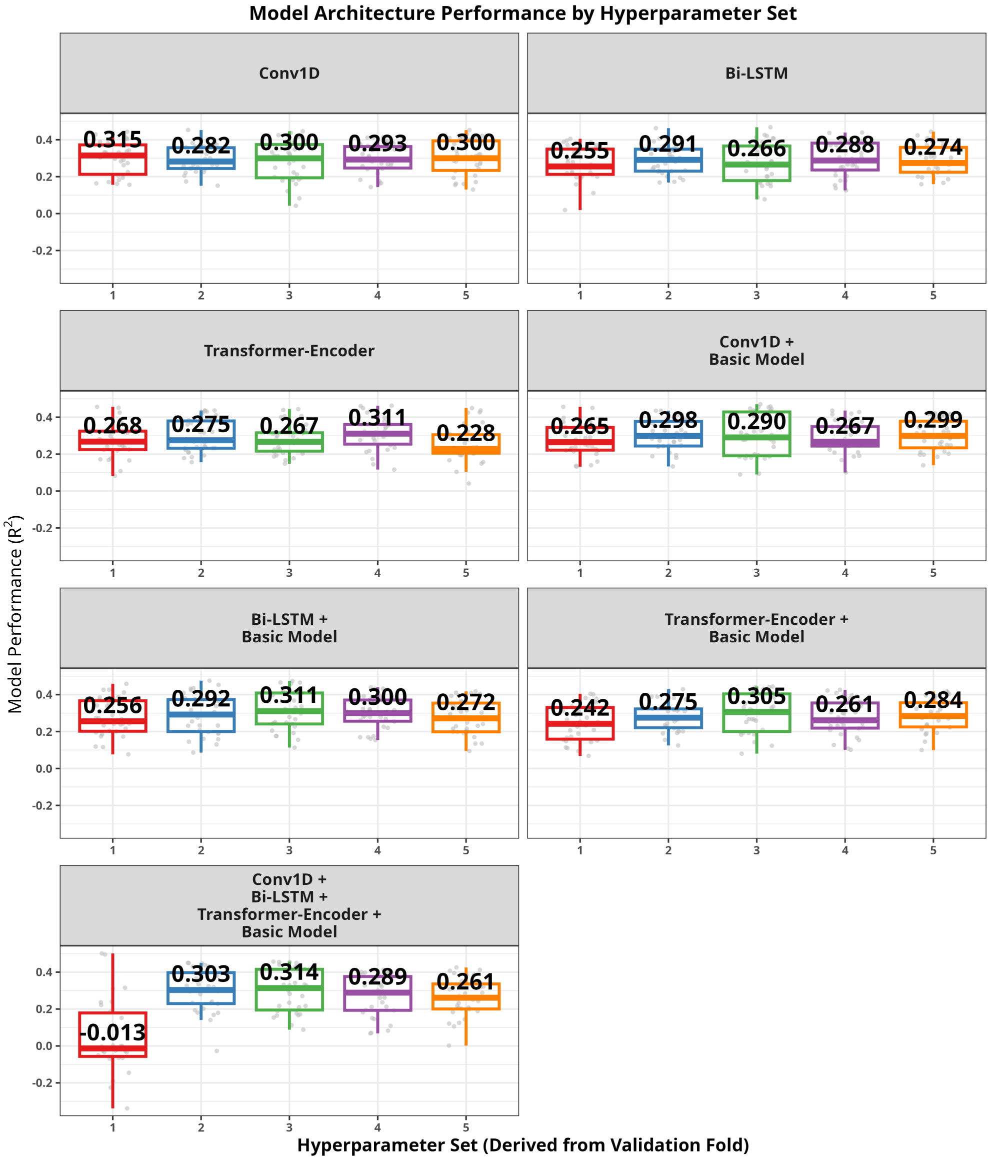


**Figure S6**: **Architecture Evaluation Split by Validation Set.** With the hyperparameter optimization scheme from Fig 1, 5 different validation datasets produced 5 different hyperparameter sets. Each graph has the median $R^2$ labeled.

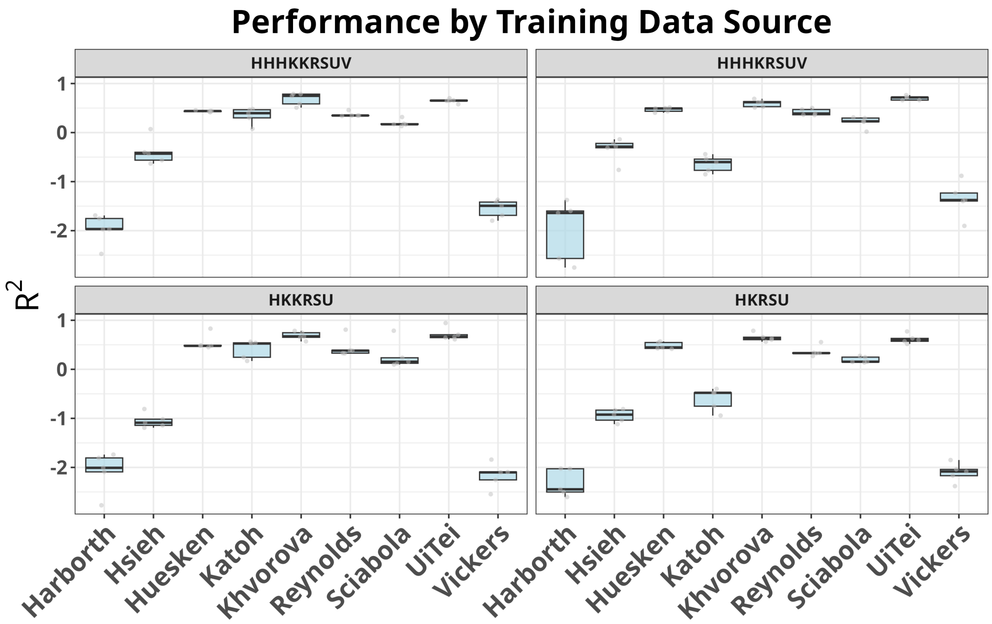


**Figure S7**: **Performance of Learned Tokenized Embedding with Different Dataset Builds.** Different combinations of datasets where used to build models based on the optimization from Fig 2D then tested for prediction performance on the individual datasets. The Huesken dataset was used with capped normalization. The different combined datasets are: HHHKKRSUV (Harborth, Hsieh, Huesken, Katoh, Khvorova, Reynolds, Sciabola, UiTei, and Vicker), HHHKRSUV (Harborth, Hsieh, Huesken, Khvorova, Reynolds, Sciabola, UiTei, and Vicker), HKKRSU (Huesken, Katoh, Khvorova, Reynolds, Sciabola, UiTei) and HKRSU (Huesken, Khvorova, Reynolds, Sciabola, UiTei).

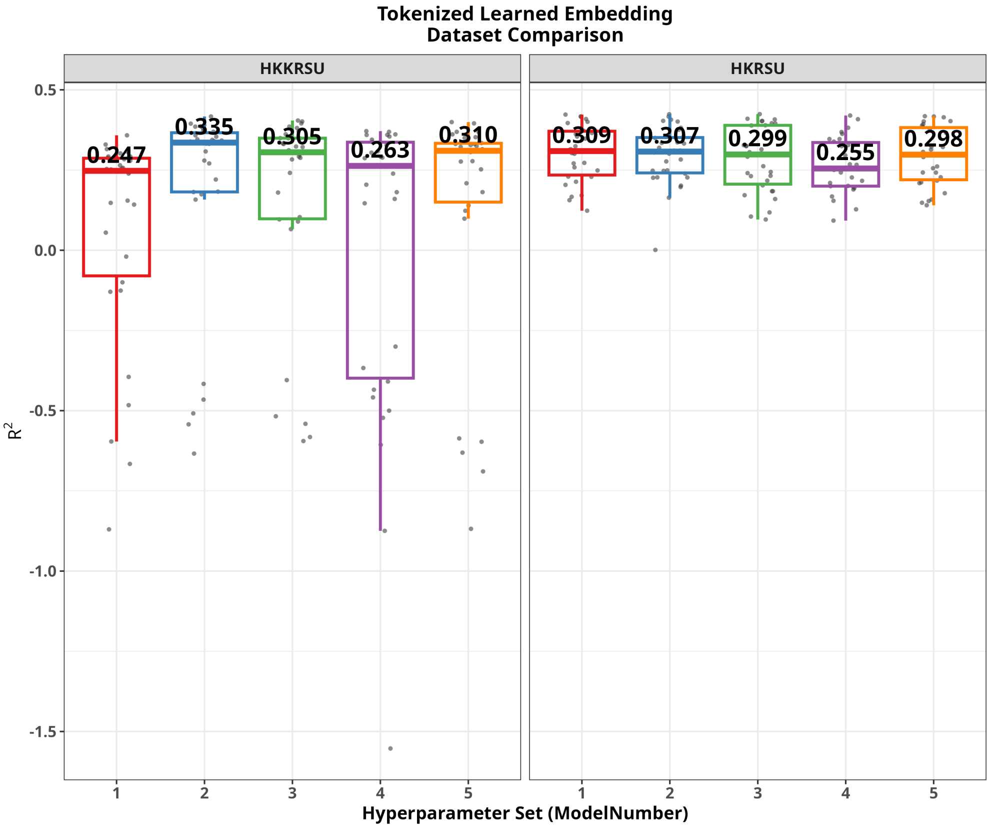


**Figure S8**: **Inclusion of Katoh dataset increases $R^2$ dispersion.** Aggregate datasets of HKKRSU (Huesken, Katoh, Khvorova, Reynolds, Sciabola, UiTei) and HKRSU (Huesken, Khvorova, Reynolds, Sciabola, UiTei), the latter not including the Katoh dataset, were evaluated with tokenized learned embedding with the evaluation methods from Fig 1. Inclusion of the Katoh dataset led to a larger spread in $R^2$ values indicating this dataset may reduce some of the predictive power on unseen data when included in the model.

**Table S1**: **Optimized tokenized learned embedding hyperparameters.** Each row represents a hyperparameter, and each column represents the set of optimized values derived from a different validation fold from 5-fold CV.

| Hyperparameter | Set 1 | Set 2 | Set 3 | Set 4 | Set 5 |
|---|---|---|---|---|---|
| **Learning rate** | 1.55e-4 | 1.00e-5 | 1.75e-4 | 8.26e-4 | 2.06e-4 |
| **siRNA Embedding Output** | 67 | 96 | 113 | 38 | 165 |
| **siRNA Dropout** | 0.0475 | 0.179 | 0.00716 | 0.0338 | 0.253 |
| **3'mRNA Flank Embedding Output** | 10 | 23 | 17 | 1 | 19 |
| **3'mRNA Flank Dropout** | 0.115 | 0.215 | 0.146 | 0.299 | 0.0751 |
| **5'mRNA Flank Embedding Output** | 16 | 3 | 15 | 215 | 6 |
| **5'mRNA Flank Dropout** | 0.255 | 0.246 | 0.286 | 0.287 | 0.0161 |
| **FFN layers** | 2 | 3 | 8 | 6 | 2 |

| Set | Layer | Activation | Output Units | Dropout |
|---|---|---|---|---|
| **Set 1** | Layer 1 | mish | 206 | 0.174 |
| | Layer 2 | relu | 119 | 0.194 |
| **Set 2** | Layer 1 | mish | 258 | 0.182 |
| | Layer 2 | mish | 113 | 0.205 |
| | Layer 3 | relu | 267 | 0.185 |
| **Set 3** | Layer 1 | relu | 134 | 0.151 |
| | Layer 2 | tanh | 365 | 0.205 |
| | Layer 3 | elu | 351 | 0.239 |
| | Layer 4 | linear | 131 | 0.224 |
| | Layer 5 | relu | 440 | 0.0607 |
| | Layer 6 | mish | 331 | 0.131 |
| | Layer 7 | relu | 425 | 0.167 |
| | Layer 8 | relu | 360 | 0.0542 |
| **Set 4** | Layer 1 | elu | 313 | 0.280 |
| | Layer 2 | mish | 479 | 0.285 |
| | Layer 3 | elu | 388 | 0.169 |
| | Layer 4 | tanh | 243 | 0.116 |
| | Layer 5 | mish | 334 | 0.255 |
| | Layer 6 | elu | 180 | 0.00231 |
| **Set 5** | Layer 1 | linear | 327 | 0.127 |
| | Layer 2 | elu | 322 | 0.171 |
